# How to entrain a selected neuronal rhythm but not others: open-loop dithered brain stimulation for selective entrainment

**DOI:** 10.1101/2022.07.06.499051

**Authors:** Benoit Duchet, James J. Sermon, Gihan Weerasinghe, Timothy Denison, Rafal Bogacz

## Abstract

While brain stimulation therapies such as deep brain stimulation for Parkinson’s disease can be effective, they have yet to reach their full potential across neurological disorders. Entraining neuronal rhythms using rhythmic brain stimulation has been suggested as a new therapeutic mechanism to restore neurotypical behavior in conditions such as chronic pain, depression, and Alzheimer’s disease. However, theoretical and experimental evidence indicate that brain stimulation can also entrain neuronal rhythms at sub- and super-harmonics, far from the stimulation frequency. Crucially, these counterintuitive effects can be harmful to patients, for example by triggering debilitating involuntary movements in Parkinson’s disease. We therefore propose a principled approach to selectively promote rhythms close to the stimulation frequency, while avoiding potential harmful effects by preventing entrainment at sub- and super-harmonics. Our open-loop approach to selective entrainment, dithered stimulation, consists in adding white noise to the stimulation period. We theoretically establish the ability of dithered stimulation to selectively entrain a given brain rhythm, and verify its efficacy in simulations of uncoupled neural oscillators, and networks of coupled neural oscillators. Furthermore, we show that dithered stimulation can be implemented in neurostimulators with limited capabilities by toggling within a finite set of stimulation frequencies. Likely implementable across a variety of existing brain stimulation devices, dithering-based selective entrainment has potential to enable new brain stimulation therapies, as well as new neuroscientific research exploiting its ability to modulate higher-order entrainment.

## 1 Introduction

In humans, neuronal rhythms can be entrained non-invasively using periodic stimuli such as auditory stimulation [1, 2], visual stimulation [3, 4, 5, 6], or transcranial stimulation [7, 8, 9, 10, 11, 12]. Animal studies demonstrate that transcranial electrical stimulation can reliably entrain individual cortical neurons [13, 14], and can even entrain neurons in the hippocampus and the basal ganglia [15]. Additionally, rhythmic sensory stimulation in humans provides evidence for the entrainment of neural oscillators by stimulation over a simple sequence of evoked responses [4, 2]. Deep brain stimulation (DBS), which invasively delivers electrical stimulation to deep targets in the brain, was also shown to entrain basal ganglia neurons in humans [16].

In line with this evidence, entraining neuronal rhythms using brain stimulation has been suggested as a new therapeutic mechanism to restore neurotypical behavior. Entraining individual alpha rhythms (8-12 Hz) using transcranial stimulation shows promise in patients with depression [17, 11] and chronic pain [18]. Gamma frequency (30-100 Hz) entrainment attenuates pathology associated with Alzheimer’s disease and improves hippocampal function in mice [19, 20]. It was recently shown that gamma entrainment may also favorably influence cognitive function as well as biomarkers of Alzheimer’s-disease-associated degeneration in humans [21]. In patients with Parkinson’s disease (PD), low-frequency switching of DBS between hemispheres entrains stepping, and could in principle be used to ameliorate gait impairment [22]. Transcranial alternating current stimulation at gamma frequency improves movement velocity in PD patients [23], likely by entraining the prokinetic gamma rhythm.

However, neuronal rhythms far from the stimulation frequency can also be inadvertently entrained by periodic brain stimulation, which, crucially, may lead to harmful effects. In patients with PD, finely-tuned gamma oscillations [24] can be entrained at half the frequency of DBS (see Fig 1A), which may be linked to debilitating involuntary movements known as dyskinesia [25, 26, 27]. In a study involving a canine with epilepsy, the frequency of DBS was chosen to avoid sub-harmonic entrainment of rhythms associated with epileptic seizures [28]. Furthermore, sensory stimulation using visual flashes at 10 Hz can lead to super-harmonic entrainment [3], and was also reported to cause undesirable side effects as highlighted in a recent commentary [29].

**Figure 1:**
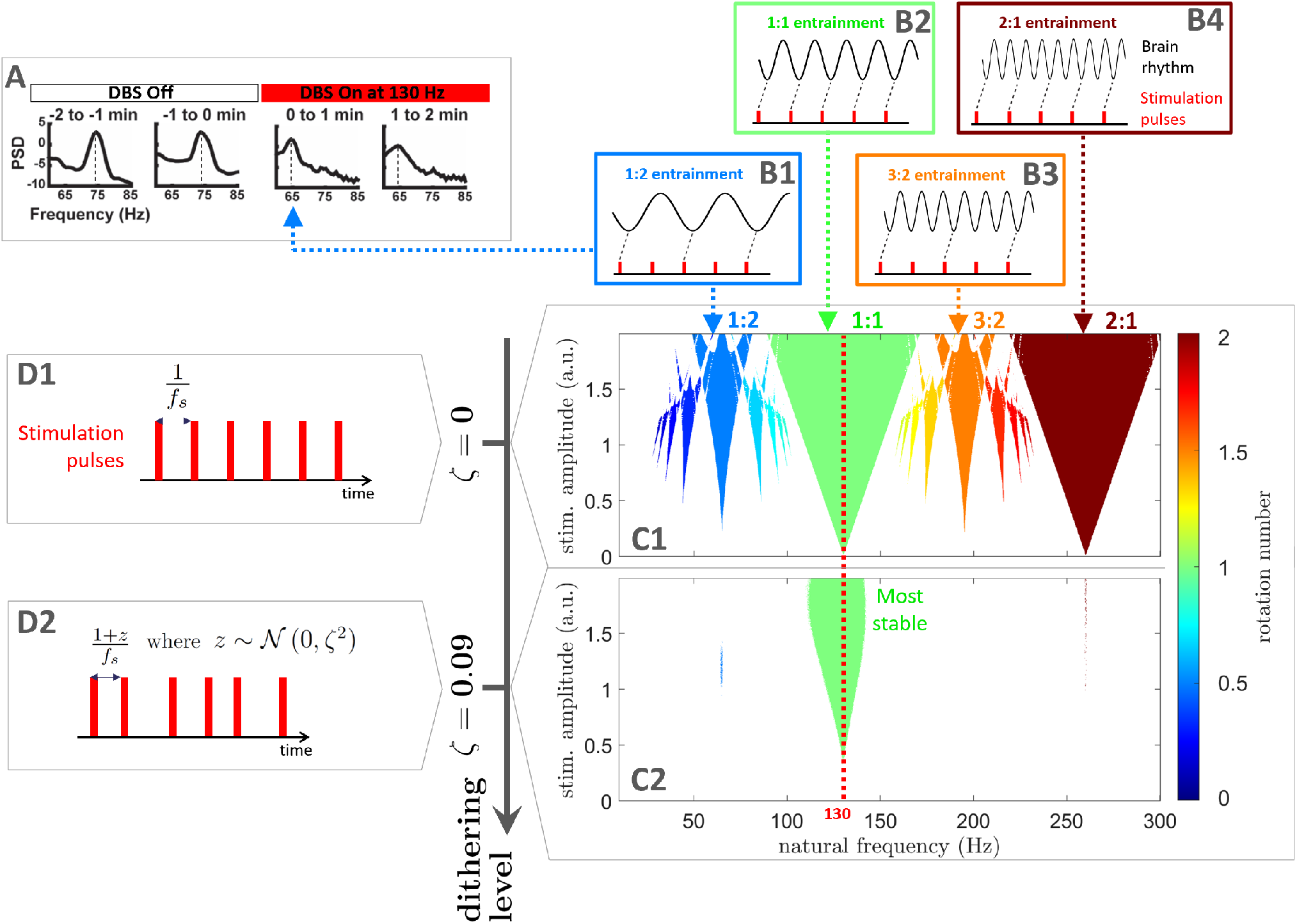
Selective entrainment of brain rhythms using dithered stimulation. When stimulation is perfectly periodic as depicted in **D1** (*f*_*s*_ denotes the stimulation frequency), neural oscillators may be entrained at the stimulation frequency but also at sub- and supra-harmonics of the stimulation frequency. Corresponding entrainment regions (Arnold tongues) are represented in **C1** for uncoupled neural oscillators modelled using the sine circle map (described in Section 2.1). Stimulation is provided at 130 Hz (red dashed line), with stimulation amplitude shown on the vertical axis, and the natural frequency of oscillators on the horizontal axis. Entrainment is observed when the rotation number (color scale in **C**) is an integer ratio (only regions of frequency-locking determined as described in Section 4.2 are shown). Entrainment at various rotation numbers is illustrated in **B1-B4**. As an example, DBS in PD patients can entrain cortical gamma oscillations at half the stimulation frequency, which corresponds to 1:2 entrainment (**A**, adapted from [25] with no permission required). With dithering, stimulation is not perfectly periodic (**D2**), and past a certain level of noise in the stimulation period, only the 1:1 Arnold tongue subsists (green tongue in **C2**).

Synchronisation theory predicts that rhythms that are close to sub- and super-harmonics of the stimulation frequency may be entrained by stimulation [30]. Neural oscillators can be assumed to have a natural frequency at which they tend to oscillate. Under certain conditions, and if stimulation is strong enough, the oscillation frequency can be shifted towards the stimulation frequency or its harmonics (more details in Section 2.1). In neural networks, the possibility of sub- and super-harmonic entrainment has been corroborated by computational models [31, 32, 33, 34, 35, 27]. The frequency locking behaviour of a rhythm to external stimulation is characterised by its rotation number, which corresponds to the average number of cycles achieved by the rhythm between two periodic stimulation pulses. When the rotation number is rational, i.e. of the form *p*:*q* with *p* and *q* coprime integers, the rhythm is entrained by stimulation (see examples in Fig 1B1-4). Synchronisation regions in the stimulation frequency/amplitude space form characteristic patterns called Arnold tongues [36] (see examples in Fig 1C1). Arnold tongues at all possible integer ratios are predicted for non-linear systems close to Hopf bifurcations [37]. However, among higher-order entrainment ratios (*p*:*q* with *p* > 1 and *q* > 1), only the most stable ones (low *p* and *q*) are likely to be observed in real neural systems. In keeping with this, experimental evidence of higher-order entrainment to brain stimulation is so far limited to the most stable higher-order ratios such as 1:2, 2:1, 3:1, and 4:1 [3, 38, 25, 26].

To promote a target physiological rhythm using brain stimulation while avoiding harmful effects, it would therefore be desirable to entrain the target neuronal rhythm while ensuring that other rhythms are not entrained by stimulation. We call such a strategy “selective entrainment”. In this study, we propose a simple method to achieve selective entrainment, which we call “dithered stimulation”. This method is open-loop, and consists in introducing noise in the stimulation period. We present a theoretical basis for the efficacy of dithered stimulation, and confirm its effectiveness in computational models representing uncoupled neural oscillators, and populations of coupled neural oscillators. Additionally, we describe how our dithering approach could be implemented in existing neurostimulators.

## 2 Results

Our approach to selective entrainment, dithered stimulation, rests on the fact that entrainment is the most stable around the stimulation frequency. In other words, 1:1 entrainment is generally more stable than *p*:*q* entrainment for *p* > 1 and *q* > 1. Because the 1:1 tongue is generally larger, small changes in oscillator frequency do not affect the 1:1 Arnold tongue much, while higher-order Arnold tongues are less stable to perturbations. This is also true for small changes in stimulation frequency. Therefore, introducing variations in the stimulation frequency should perturb frequency locking more for higher-order tongues than for 1:1 entrainment. This is the basis of our dithering approach to selective entrainment, which consists in its simplest form in adding white noise to the stimulation period, as illustrated in Fig 1D2. Open loop stimulation patterns with irregular pulse timings have been investigated for other purposes [39, 40, 41, 42]. Here, we consider an open loop stimulation pattern where the time interval between stimulation pulses always changes and is given by (1 + *z*)/*f*_*s*_, where *f*_*s*_ is the base stimulation frequency, and *z* is a normal random number sampled from 𝒩 (0, *ζ*^2^) for each stimulation interval. We call the standard deviation *ζ* “dithering level”. Past a certain dithering level, only the 1:1 Arnold tongue remains (Fig 1C2), which ensures that only neuronal rhythms of frequency close to the stimulation frequency are entrained. By adjusting *f*_*s*_ to the target rhythm to entrain, selective entrainment can therefore be achieved using dithered stimulation. We next provide theoretical and computational demonstrations of the efficacy of the method. Specifically, we will first consider a simple model of uncoupled neural oscillators (the sine circle map) for which it is possible to analytically approximate the width of Arnold tongues. We will then show that the conclusions generalize to a more complex model often used to describe neural oscillations (the Kuramoto model).

### 2.1 Selective entrainment can be achieved by dithered stimulation in models of uncoupled neural oscillators

The sine circle map is the simplest model describing the influence of periodic stimulation on a single neural oscillator, and can be used to provide a theoretical basis for the efficacy of dithered stimulation as a selective entrainment strategy. The model maps the phase of an oscillator right before stimulation pulse *n* (denoted *θ*_*n*_) to the phase of the oscillator right before stimulation pulse *n* + 1 (denoted *θ*_*n*+1_). The map can be written as

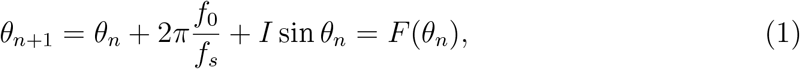

where *f*_0_ is the oscillator natural frequency, *f*_*s*_ the stimulation frequency, and *I* the stimulation magnitude. Entrainment can arise because a stimulus may advance or delay the phase of an oscillator depending on the phase at which it is applied. This concept is captured by the phase response curve (PRC) of the oscillator, which describes the change in phase of the oscillator as a function of the stimulation phase. The PRC of the sine circle map is a simple sinusoid (*Z*(*θ*) = sin *θ*). Since brain oscillations can manifest across a wide range of frequencies, we consider a population of uncoupled neural oscillators modelled by equation (1) where *f*_0_ corresponds to the natural frequency axis in Fig 1C1. For perfectly periodic stimulation, Arnold tongues are observed in Fig 1C1 at all possible entrainment ratios (rotation number obtained as detailed in Section 4.2 in the Methods Section). When representing the rotation number as a function of natural frequency, frequency locking corresponds to plateaux where the rotation number takes a constant integer ratio across a range of natural frequencies as illustrated in Fig 2 for *ζ* = 0 (only ratios with low *p* and *q* are easily discernable).

**Figure 2:**
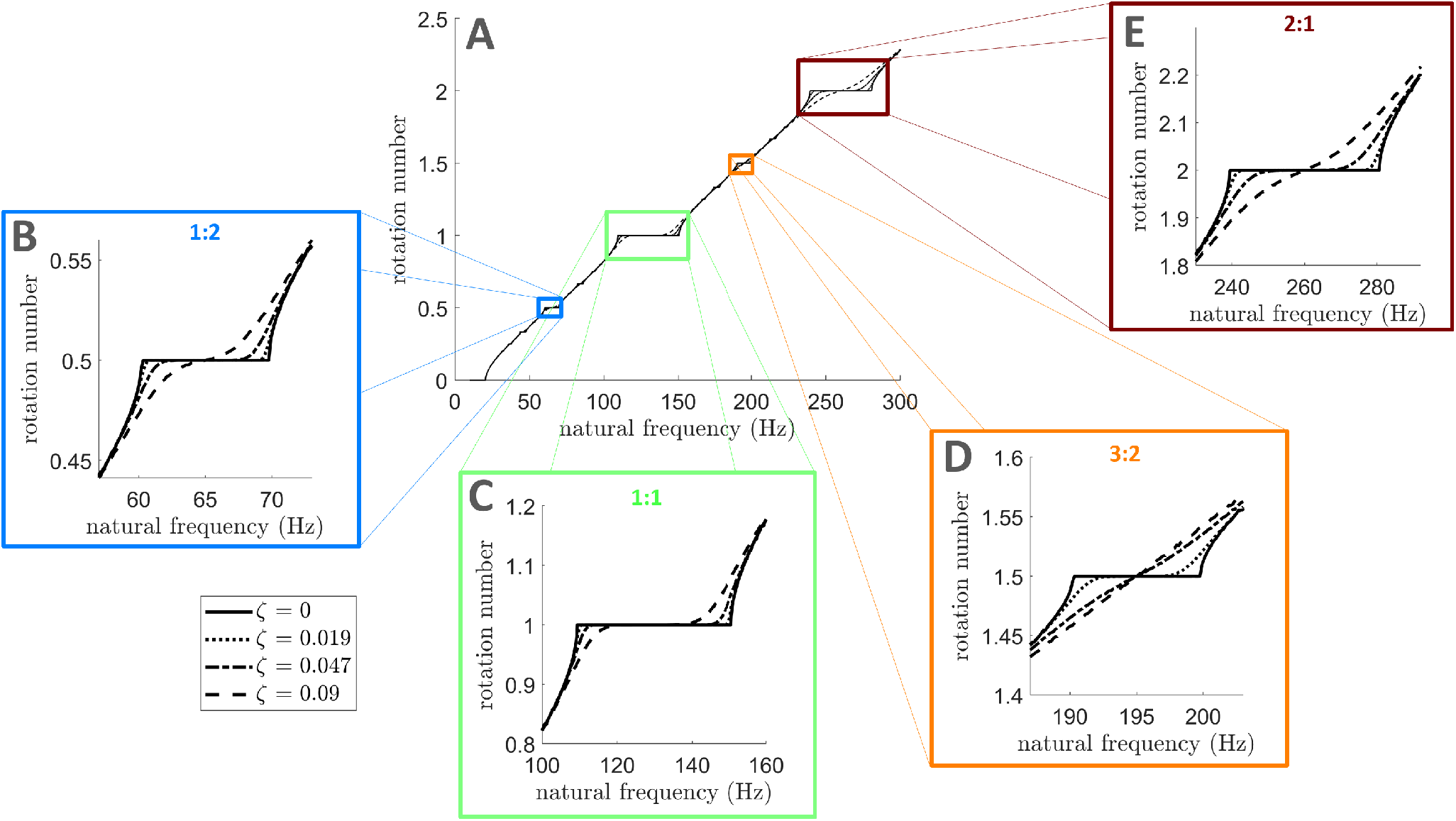
Frequency locking plateaux disappear at lower dithering levels for higher-order entrainment than for 1:1 entrainment. Rotation number in the sine circle map as a function of oscillator frequency, for *f*_*s*_ = 130 Hz, and *I* = 1. Solid lines correspond to perfectly periodic stimulation, while dashed/dotted lines correspond to dithered stimulation with increasing dithering levels. While a large part of the 1:1 frequency locking plateau is preserved for all dithering levels shown (**C**), plateaux for 1:2 (**B**), 3:2 (**D**), and 2:1 (**E**) frequency locking have disappeared at *ζ* = 0.09. Panels **B-E** are zoomed-in versions of panel **A**. For each dithering level, the rotation number is averaged over 10 repeats, with 10^4^ stimulation pulses per repeat.

#### 2.1.1 Theoretical justification of dithering stimulation as a selective entrainment strategy

To demonstrate analytically that dithered stimulation destabilises the most prominent higher-order entrainment ratios more than 1:1 entrainment, we introduce dithered stimulation in the sine circle map. The sine circle map with dithered stimulation becomes the stochastic map

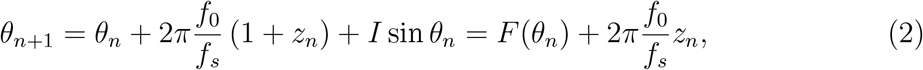

where the stimulation period *T*_*s*_ = 1/*f*_*s*_ is multiplied by (1 + *z*_*n*_) to model dithered stimulation, with *z*_*n*_ normal random numbers sampled from 𝒩 (0, *ζ*^2^).

We build on ideas presented in [43] to show that, as more dithering is introduced in equation (2), the relative decrease in width of the most prominent higher-order Arnold tongues is greater than that of the 1:1 tongue. The most prominent higher-order tongues are of the form *p*:1 with *p* > 1 (supra-harmonic entrainment), and (2*p*−1):2 with *p* ≥ 1. In Fig 1C1, these correspond to the 2:1 tongue, and to the 1:2 and 3:2 tongues, respectively. We denote by **Δ***f*^*p*:*q*^(*I, ζ*) the range of oscillator natural frequencies that can be entrained by the *p*:*q* tongue with dithered stimulation of amplitude *I* and dithering level *ζ*, i.e. the width of the tongue at that stimulation amplitude and dithering level. Approximate expressions for Δ*f*^*p*:1^(*I, ζ*), *p* ≥ 1 and Δ*f* ^(2*p*−1):2^(*I, ζ*), *p* ≥ 1 are derived in Section 4.1 in the Methods Section assuming *ζ* ≪ 1 and *I* ≪ 1. Neglecting small high order *I* terms in equation (7) in Section 4.1, the width of *p*:1 tongues (*p* ≥ 1) with dithered stimulation relative to the width of the same tongues with perfectly periodic stimulation can be obtained as

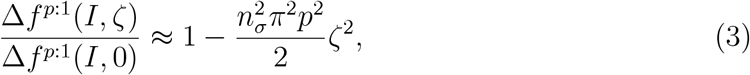

where *n*_*σ*_ quantifies the number of standard deviations of the jump distribution beyond which temporary escapes of the basin of attraction of the periodic orbit are considered to not significantly affect the locking behavior (see Section 4.1 in the Methods Section for more details). The value of *n*_*σ*_ is taken to be the same across tongues. Similarly, neglecting high order *I* terms in equation (10) in Section 4.1, the width of (2*p* − 1):2 tongues (*p* ≥ 1) with dithered stimulation relative to the width of the same tongues with perfectly periodic stimulation is

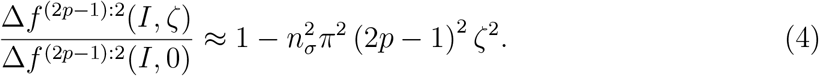

It follows that the relative decrease of the most prominent higher-order tongues (any *p* > 1 in equation (3) and any *p* ≥1 in equation (4)) with increasing dithering levels is always greater than for the 1:1 tongue (*p* = 1 in equation (3)). This result is valid for any stimulation frequency and underlies the efficacy of dithered stimulation for selective entrainment.

#### 2.1.2 Validation using simulations of uncoupled neural oscillators

To confirm that there exist a dithering level at which the 1:1 tongue displays a broad frequency locking region while other tongues have disappeared, we simulate the sine circle map with dithered stimulation (equation (2)) for increasing noise levels. As an example, we set the base stimulation frequency to *f*_*s*_ = 130 Hz, which corresponds to the frequency of clinically available DBS. As *ζ* is increased, frequency locking plateaux in Fig 2 disappear faster for higher order entrainment than for 1:1 entrainment. Significant 1:1 frequency locking is still possible for *ζ* = 0.009 as indicated by the large dashed line plateau in Fig 2C, but no 1:2, 3:2, or 2:1 frequency locking is possible (Fig 2B,D-E). Similarly, higher order Arnold tongues become narrower faster than the 1:1 tongue as *ζ* is increased (Fig 3A1-A4). For *ζ* = 0.009, the 1:1 tongue can still entrain neural oscillators with natural frequencies in the vicinity of the stimulation frequency, while other tongues have disappeared (Fig 3A4).

**Figure 3:**
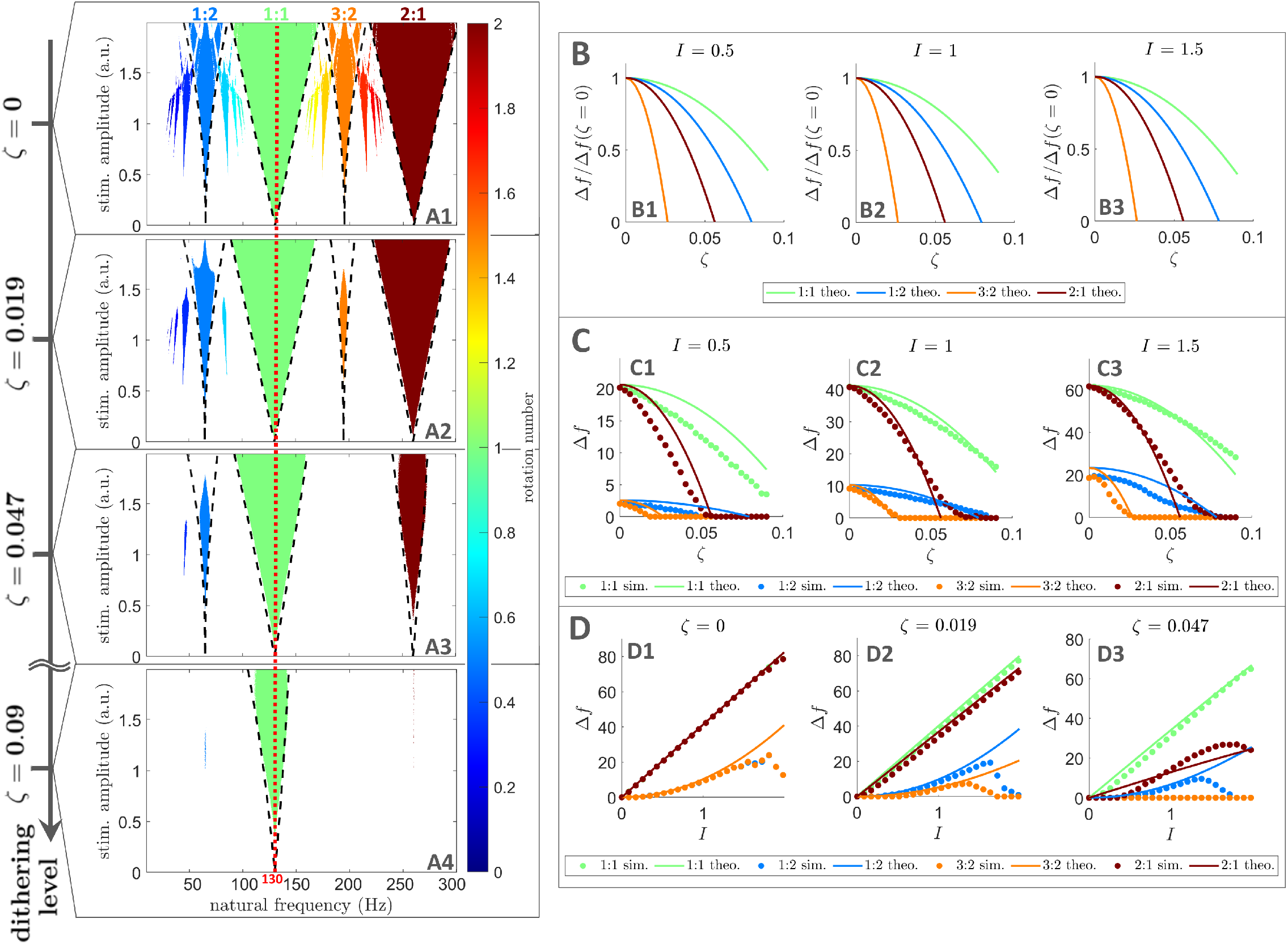
Arnold tongues disappear at lower dithering levels for higher order entrainment than for 1:1 entrainment in uncoupled neural oscillators. **A:** Frequency locking regions in the oscillator frequency/stimulation amplitude space. Only regions of frequency-locking (determined as presented in Section 4.2), known as Arnold tongues, are shown in color. The color scale represents the rotation number. Dithering level increases from top to bottom, and theoretical tongue boundaries (equations (6) and (9)) are shown by black dashed lines. The stimulation frequency is indicated by a red dashed line. An animation with finer steps in dithering levels is provided as supplementary material (S1.mp4, see S1 for caption). **B:** Showing theoretical tongue width normalised by its value for perfectly periodic stimulation, plotted against dithering level, for three stimulation amplitudes. The dependence on stimulation amplitude is very slight. **C:** Comparing tongue width obtained from theory and simulations, as a function of dithering level, for three stimulation amplitudes. **D:** Comparing tongue width obtained from theory and simulations, as a function of stimulation amplitude, for three dithering levels. In all panels showing data from simulations, for each natural frequency, stimulation amplitude, and dithering value, the rotation number is averaged over 10 repeats, with 10^4^ stimulation pulses per repeat. In panels **B-D**, theoretical tongue widths refer to equations (7) and (10).

Simulations of the sine circle map with dithered stimulation also validate our theoretical results. More specifically, equations (6) and (9) (Methods Section) describing tongue boundaries in the presence of dithered stimulation (dashed lines in Fig 3A1-A4) approximately match tongue boundaries obtained directly from simulations. Additionally, Fig 3B is consistent with the faster relative decrease in width of higher order tongues compared to the 1:1 tongue as the dithering level is increased, and tongue width measurements from simulations approximately match theoretical values from equations (7) and (10) (Methods Section) as shown in Fig 3C-D. We also confirm that our theoretical results hold for *p*:1 tongues and (2*p* − 1):2 tongues for larger values of *p* in Figs 8 and 9 in the supplementary information. As predicted, these tongues disappear for even lower dithering levels than the tongues in Fig 3. In Figs 3, 8, and 9, theoretical results were based on *n*_*σ*_ = 4, i.e. frequency-locking was considered to occurs when at least 99.99% of locking cycles do not escape the periodic orbit. This was found to robustly correspond to frequency-locking plateaux in Fig 2 across noise levels.

Although a range of dithering levels that suppress the 1:2 tongue while still preserving the 1:1 tongue were found (Fig 3A4), we note that the 1:2 tongue is the hardest higher-order tongue to destabilise (see Fig 3B2 and C2). The fact that 1:2 entrainment was the first type of higher-order entrainment reported in patients with Parkinson’s disease treated with DBS [25, 26] is in line with these predictions.

We showed that the efficacy of dithered stimulation as a selective entrainment strategy is supported by theory, and confirmed by simulations of uncoupled neural oscillators. However brain signals such as local field potentials can be better described by networks of coupled oscillators representing coupled neurons or coupled micro-circuits [44, 45, 46]. Thus, we next test dithered stimulation in this more realistic setting.

### 2.2 Selective entrainment can be achieved by dithering in populations of coupled neural oscillators

In order to test dithered stimulation as a selective entrainment strategy in a more realistic setting, we simulate populations of coupled neural oscillators using the Kuramoto model [47]. For each natural frequency *f*_0_ in the frequency range of interest, we consider a population of *M* = 100 phase oscillators with homogeneous coupling, natural frequencies distributed around *f*_0_, and the PRC of a standard Hodgkin-Huxley neuron [48, 49] (see Fig 12B1 in the supplementary information). Full details on the model can be found in Section 4.3 in the Methods Section. As opposed to the sine circle map, the Kuramoto model is a continuous-time model. Thus, we are also able to use more realistic, square stimulation pulses with a temporal extent, and a negative component for charge balance (see Section 4.3 and Fig 12B2 in the supplementary information).

As shown in Fig 4, dithered stimulation is an effective selective entrainment strategy in populations of coupled neural oscillators. For perfectly periodic stimulation at 130 Hz, populations of coupled neural oscillators can be entrained at the stimulation frequency (1:1 entrainment), but also at higher-order entrainment ratios for certain natural frequencies and stimulation amplitudes (Fig 4A1). Details on how entrainment metrics presented in Fig 4 are obtained can be found in Section 4.2 in the Methods Section. In the natural frequency range considered, the only higher-order tongues with non-zero widths are the 1:2, 3:2, and 2:1 tongues, which were identified as the most prominent higher-order tongues in Section 2.1. As the dithering level *ζ* is increased, these higher-order tongues fade, while the 1:1 tongue is mostly preserved (Fig 4A1-A4). For *ζ* = 0.15, 1:1 entrainment is maintained for a large range of natural frequencies and stimulation amplitudes, while higher order entrainment has vanished (Fig 4A4). This is confirmed by measuring the width of Arnold tongues as a function of stimulation amplitude (Fig 4B4).

**Figure 4:**
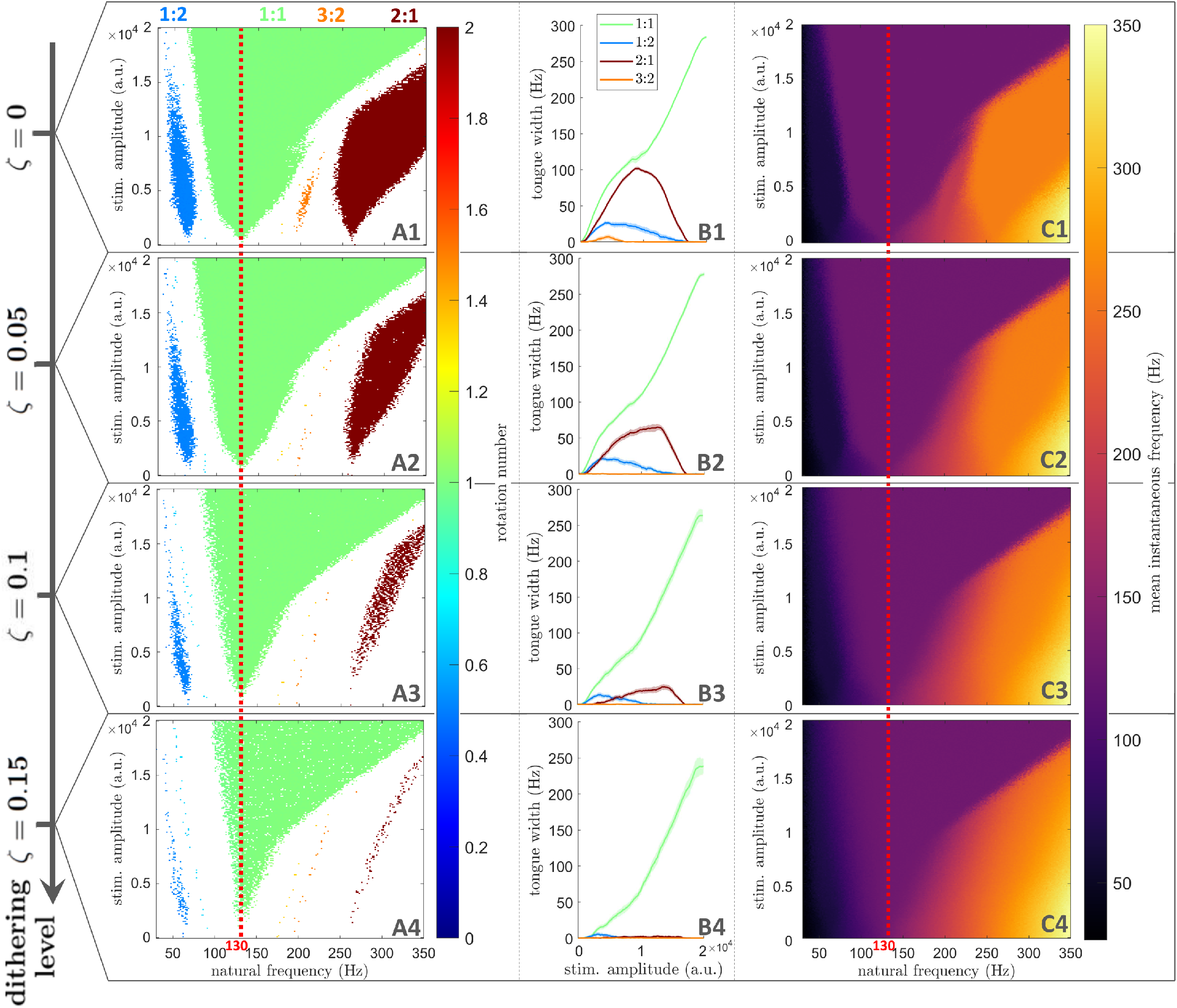
Arnold tongues disappear at lower dithering levels for higher-order entrainment than for 1:1 entrainment in populations of coupled neural oscillators. Dithering level increases from the top row to the bottom row. **A:** Frequency locking regions in the natural frequency/stimulation amplitude space. Only regions of frequency-locking (determined as presented in Section 4.2), are shown in color. The color scale represents the rotation number. **B:** Arnold tongue width as a function of stimulation amplitude. **C:** Mean instantaneous frequency (represented by the color scale) in the natural frequency/stimulation amplitude space. In **A** and **C**, for each natural frequency, stimulation amplitude, and dithering value, the rotation number or mean instantaneous frequency are averaged over 5 repeats, with 400 stimulation pulses per repeat. The stimulation frequency is indicated by a red dashed line.

The variation in the mean instantaneous frequency of the Kuramoto populations with respect to *f*_0_ also supports this conclusion. For *ζ* = 0.15, the mean instantaneous frequency is constant in the 1:1 tongue, signalling frequency-locking to the stimulation frequency, while a non-zero frequency gradient along *f*_0_ is observed elsewhere, indicated the absence of frequency-locking (Fig 4C4). This was not the case for the perfectly periodic case, where regions of constant mean instantaneous frequency can be seen to approximately match the 1:2, 3:2, and 2:1 tongues (Fig 4C1). Further validation is provided in Fig 10 (supplementary information) based on the phase locking value (PLV), which was used to assess 1:1 synchronisation for example in [6, 50] (see Section 5.1 in the supplementary information for more details).

### 2.3 Dithering can be implemented by toggling within a finite set of stimulation frequencies

To ensure that dithered stimulation can be implemented in a broad range of existing neurostimulators, we consider different ways of toggling within a finite set of stimulation frequencies as approximations of white noise based dithered stimulation. Let us consider a set of *n* stimulation frequencies *S*_*n*_ = {*f*_*s*,1_, *f*_*s*,2_, …, *f*_*s,n*_}, such that {1/*f*_*s*,1_, 1/*f*_*s*,2_, …, 1/*f*_*s,n*_} are symmetrically distributed around 1/*f*_*s*_, where *f*_*s*_ is the base stimulation frequency. The simplest way of approximating white noise based dithered stimulation is to randomly select a stimulation frequency from the set *S*_*n*_ for each stimulation period. This random cycling approach is illustrated in Fig 5A. If the neurostimulator is unable to generate random numbers, deterministic cycling can be implemented by toggling from one stimulation frequency in the set to the next (i.e. from *f*_*s,i*_ to *f*_*s,i*+1_ for *i* = {1, ‥, *n* − 1}, and from *f*_*s,n*_ to *f*_*s*,1_) at each stimulation period (Fig 5B). If the device is unable to toggle between frequencies at each stimulation period, toggling can be slower and take place only after *N*_*r*_ stimulation periods at the same stimulation frequency. This slow deterministic cycling is shown for *N*_*r*_ = 2 in Fig 5C.

**Figure 5:**
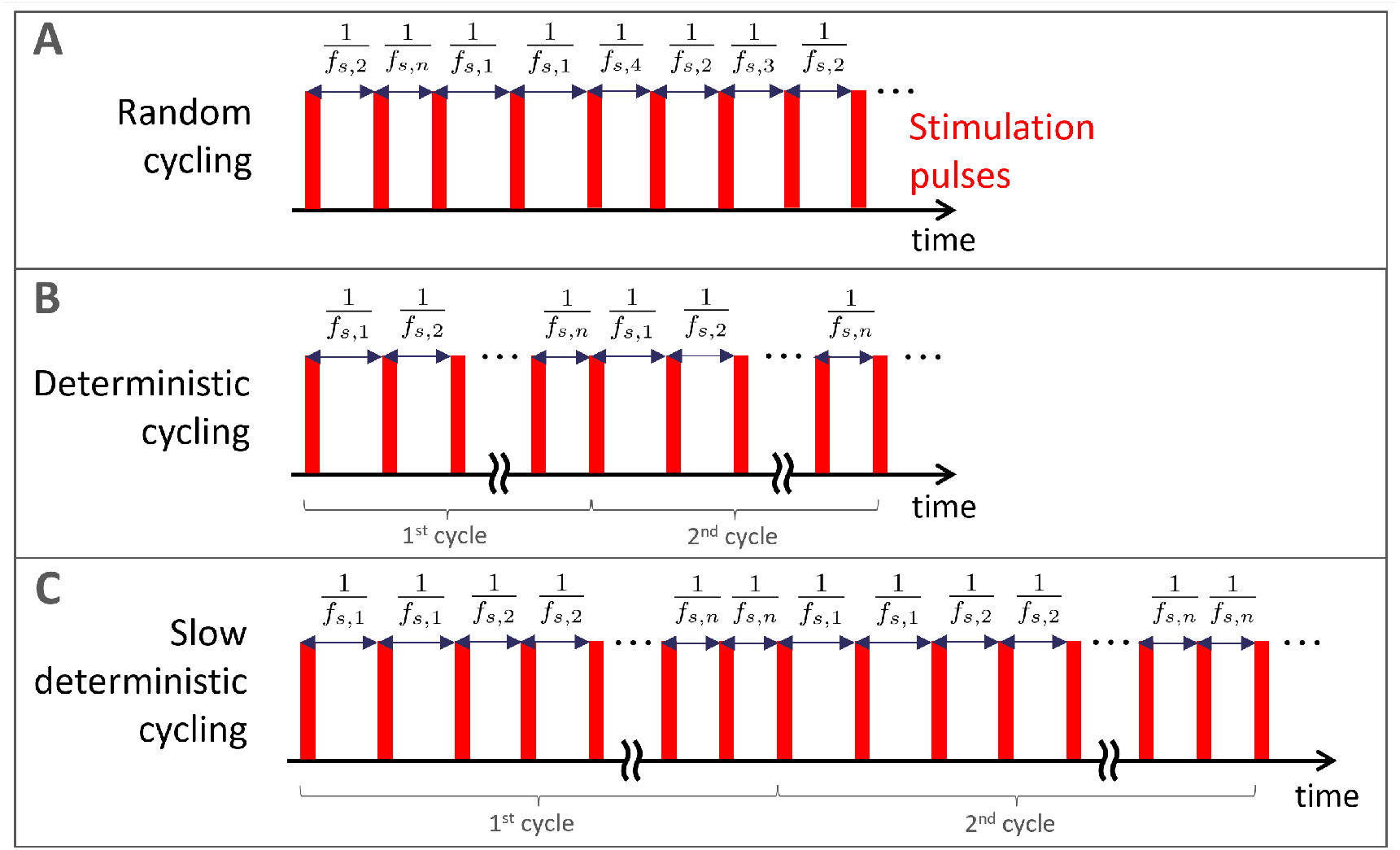
Pulse trains approximating noise based dithered stimulation by toggling within a finite set of stimulation frequencies. **A:** Random cycling, where a stimulation frequency is selected at random for each stimulation period from a finite set of frequencies. **B:** Deterministic cycling, where stimulation frequency switch to the next frequency in the set at each stimulation period. **C:** Slow deterministic cycling, where toggling is slower and takes place only after *N*_*r*_ stimulation periods at the same stimulation frequency (*N*_*r*_ = 2 in this example). Exemplar, short stimulation pulses are shown in red, but these approaches can be used for any waveform. Panels **A** and **C** present two full cycles of the corresponding deterministic patterns.

Given a large enough set of stimulation frequencies, these toggling approaches can achieve selective entrainment in populations of coupled neural oscillators. For simplicity, the distribution of stimulation periods we pick in this analysis is uniform, and the frequency sets used are detailed in Fig 6. With only three stimulation frequencies, both random and deterministic cycling fail to realize selective entrainment (first and second rows of Fig 6). With a set of seven stimulation frequencies, higher-order tongues have vanished with both random and deterministic cycling (Fig 6A3, B3, and A4, B4). Fine structures are still visible at high frequency for deterministic cycling when plotting the mean instantaneous network frequency in the natural frequency/stimulation amplitude space (Fig 6C4). This is much less the case with this frequency set for random cycling (Fig 6C4), making random cycling preferable for a set of seven stimulation frequencies. However, with a set of 13 stimulation frequencies, this is no longer an issue for deterministic cycling (second-to-last row of Fig 6), and even slow deterministic cycling with *N*_*r*_ = 3 is effective (last row of Fig 6). These results are confirmed by PLV analysis in Fig 11 in the supplementary information.

**Figure 6:**
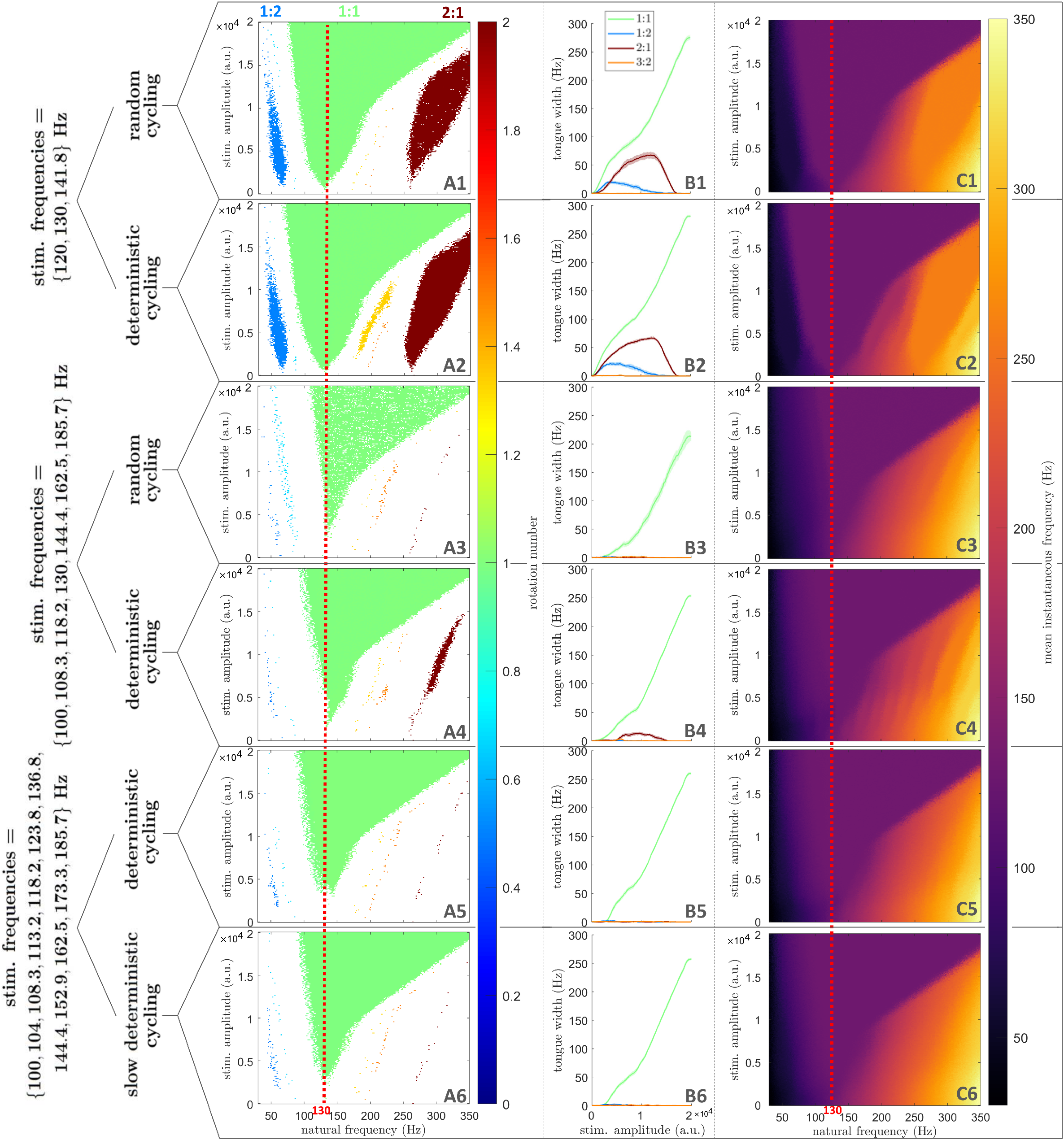
Selective entrainment can be achieved in populations of coupled neural oscillators by toggling within a finite set of stimulation frequencies. Each row corresponds to a particular type of pulse train, as indicated on the left side of the figure. For slow deterministic cycling (last row), *N*_*r*_ = 3. **A:** Frequency locking regions in the natural frequency/stimulation amplitude space. Only regions of frequency-locking (determined as presented in Section 4.2), are shown in color. The color scale represents the rotation number. **B:** Arnold tongue width as a function of stimulation amplitude. **C:** Mean instantaneous frequency (represented by the color scale) in the natural frequency/stimulation amplitude space. In **A** and **C**, for each natural frequency, stimulation amplitude, and dithering value, the rotation number or mean instantaneous frequency are averaged over 5 repeats, with 400 stimulation pulses per repeat. The stimulation frequency is indicated by a red dashed line.

In populations of coupled neural oscillators, random cycling can achieve selective entrainment with the fewest number of stimulation frequencies. Nevertheless, for devices with more limited capabilities, deterministic cycling and even slow deterministic cycling are effective when a broader set of stimulation frequencies is used.

## 3 Discussion

While entraining neural rhythms at the stimulation frequency is a promising therapy for neurological disorders, brain rhythms in different frequency bands can also be inadvertently entrained at sub- and super-harmonics of the stimulation frequency. In this study, we proposed a method to selectively entrain a given neural rhythm at the stimulation frequency, while minimising any sub- or super-harmonic entrainment that might occur in other frequency bands. Our method, which we call dithered stimulation, consists in slightly varying the stimulation period using white noise. We justified theoretically the efficacy of dithered stimulation as a selective entrainment strategy for any stimulation frequency. This was done by demonstrating analytically that the most prominent higher-order Arnold tongues shrink faster than the 1:1 Arnold tongue as the level of noise is increased in a model of uncoupled neural oscillators. The ability of dithered stimulation to selectively entrain a given rhythm was confirmed by simulations, and validated in more realistic population models of coupled neural oscillators. Additionally, we showed that dithered stimulation can be implemented in neurostimulators with limited capabilities by toggling within a finite set of stimulation frequencies, even if toggling happens on a slower time scale than the stimulation period.

### Limitations

The theoretical proof provided in Section 2.1.1 is limited to the most prominent families of higher-order Arnold tongues (*p*:1 tongues for *p* > 1 and (2*p* − 1):2 tongues for *p* ≥ 1), and to sinusoidal PRCs. Alternative theoretical investigations of the stochastic sine circle have lead to approximations of the probability density of phases [51, 52] but are not directly applicable, and it is unclear whether these approaches could yield more general analytical results. Nevertheless, our simulations show that higher-order tongues not belonging to the tongue families we theoretically considered vanish even more quickly with increasing dithering (see Fig 3A and Fig 8 in the supplementary information). Additionally, Section 2.2 demonstrates that dithered stimulation can be effective for non-sinusoidal PRCs, such as the PRC of the standard Hodgkin-Huxley neuron model (Fig 12B1 in the supplementary information), as well as when the amplitude of stimulation is not small. In general, neural oscillators and micro-circuits are expected to have a PRC with a dominant first harmonic [49, 53, 54], and our theoretical results should approximately hold.

While the sine circle map models individual neural oscillators with no amplitude variable (only a phase), the populations of coupled neural oscillators used in Section 2.2 can reproduce brain signals at the level of neural populations [44, 45, 46]. Simulations presented in Section 2.2 therefore strongly support the efficacy of dithered stimulation in neural networks. However, synaptic plasticity between coupled neural oscillators was not included because the many simulations required to cover the natural frequency/stimulation amplitude space would have been too computationally costly. Recent mean-field approximations of coupled oscillator networks with spike-timing-dependent plasticity [55] could be considered in future work to reduce computational time.

### Implications and perspectives

To date, neurostimulators are programmed without the awareness that rhythms close to integer ratios of the stimulation frequency can be inadvertently entrained. We recently proposed a therapy aimed at reinforcing neurotypical rhythms in epilepsy while being mindful of the potential entrainment effects of stimulation on seizure frequencies [28]. However this was only done by inspection, and there is a need for principled methods. For neurological disorders where healthy rhythms, pathological rhythms and/or rhythms leading to side-effects can be identified, selective entrainment based on dithering is poised to provide a robust way to reinforce healthy rhythms while avoiding undesirable effects. In case multiple target rhythms have to be selectively entrained, superimposing dithered stimulation pulse trains with different base stimulation frequencies could be investigated.

Compared to clinically available DBS, for example, the only extra parameter for dithered stimulation is the dithering level. We note that a higher dithering level was required to achieve selective entrainment in populations of coupled neural oscillators than in uncoupled neural oscillators (*ζ* = 0.15 vs *ζ* = 0.09, cf Fig 3 and Fig 4). Variations in the optimal dithering level is therefore expected depending on the target neural circuit, thus the dithering level should be adjusted experimentally. Moreover, DBS electrodes with multiple independently controlled stimulation contacts are becoming available [56, 57]. Providing dithered stimulation with different noise realisations (or different cycling patterns, if cycling through a set of stimulation frequencies) at different spatial locations within a target neural structure is likely to lower the dithering level required to achieve selective entrainment.

Beyond avoiding potentially harmful side-effects while promoting physiological rhythms, dithered stimulation offers the unique ability to modulate sub- and super-harmonic entrainment while sustaining 1:1 entrainment by modulating the dithering level. This could be used as a tool to causally probe the link between sub- or super-harmonic entrainment and behavior, such as the relationship between 1:2 entrainment of cortical gamma oscillations and dyskinesia [25, 26], or the hypothesis that 1:2 entrainment of cortical gamma oscillations by DBS promotes motor symptom alleviation [58].

### Conclusion

Selective entrainment based on dithering has potential to enable new brain stimulation therapies where there are physiological rhythms to reinforce and pathological rhythms that should not be entrained, as well as to enable new neuroscientific research. As a simple open loop stimulation strategy, it is likely to be implementable across a large variety of existing brain stimulation devices.

## 4 Methods

In this section, we approximate analytically the width of Arnold tongue families in the presence of dithered stimulation, and provide methodological details pertaining to entrainment metrics and to the simulation of networks of coupled neural oscillators.

### 4.1 Theoretical basis for the efficacy of dithered stimulation as a selective entrainment strategy

By deriving approximate expressions for the width of the most prominent families of Arnold tongues as a function of the dithering level, we propose a theoretical basis for the efficacy of dithered stimulation as a selective entrainment strategy.

#### 4.1.1 Influence of dithered stimulation on *p*:1 Arnold tongues

We consider cobweb plots to study the *p*:1 frequency locking behavior of the stochastic sine circle map given by equation (2), for *p* ≥ 1. In the deterministic case, *p*:1 frequency locking corresponds to the stable fixed point of *θ*_*n*+1_ = *F* (*θ*_*n*_) and *θ*_*n*+1_ = *θ*_*n*_ + 2*pπ*, i.e. when we have *p* complete rotations of the oscillator for every stimulation cycle. The cobweb plot representing *θ*_*n*+1_ as a function of *θ*_*n*_ has one stable and one unstable fixed points (see Fig 7A for the 1:1 case and Fig 7B for the 1:2 case). In the stochastic case, there will still be *p*:1 frequency locking if it is highly unlikely for the phase to escape the attraction “trap” between the stable and unstable fixed points in one random jump [43] (i.e. after one iteration of the stochastic map). We denote by *h* the size of the trap as indicated in Fig 7A-B.

**Figure 7:**
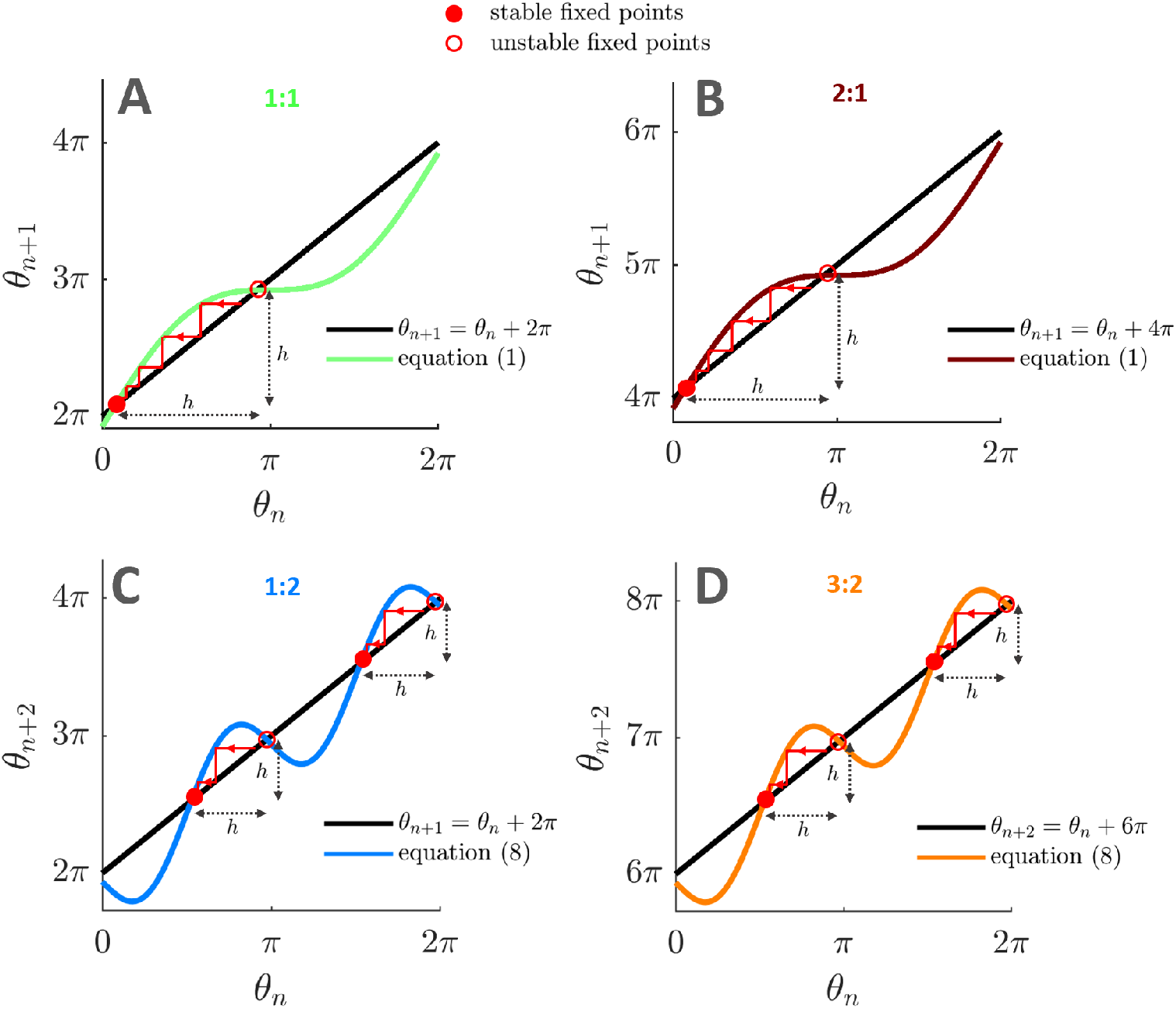
Cobweb plots of the deterministic sine circle map for different frequency locking ratios. Cobweb plots used to study frequency-locking in the deterministic sine circle map reveal the size of the trap (denoted by *h*, shortest distance between a stable and an unstable fixed point) that can prevent dithered stimulation from breaking frequency locking. Stable fixed points are identified by filled red circles, unstable fixed points by empty red circles. Parameters used are *f*_0_ = 125 Hz, *I* = 1 (A), *f*_0_ = 255 Hz, *I* = 1 (B), *f*_0_ = 63 Hz, *I* = 1.5 (C), *f*_0_ = 193 Hz, *I* = 1.5 (D). In all panels, *f*_*s*_ = 130 Hz.

The random jump size (mod 2*π*) between stimulation pulse *n* and *n* + 1 is *θ*_*n*+1_ −*θ*_*n*_ ≈ 2*π*(*f*_0_/*f*_*s*_)*z*_*n*_ since *θ*_*n*_ ≈ *F* (*θ*_*n*_) mod 2*π* in the vicinity of the stable fixed point. Therefore the jump size is approximately distributed according to 𝒩 (0, *σ*^2^) with *σ* = 2*π*(*f*_0_/*f*_*s*_)*ζ*. We consider that it is highly unlikely for the phase to clear the trap if the trap size *h* is larger than *n*_*σ*_ standard deviations of the jump size distribution, with *n*_*σ*_ ≥ 3. In [43], *n*_*σ*_ = 3, but we will consider *n*_*σ*_ ≥ 3 for the sake of generality. Therefore, frequency locking is lost when

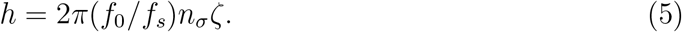

The trap size *h* can be obtained by solving for the positions of the stable and unstable fixed points of the deterministic map, and selecting the shortest absolute distance between the fixed points. We obtain for the *p*:1 case

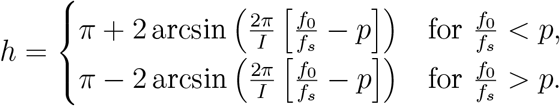

Using this result and equation (5), we obtain the *p*:1 tongue boundaries of the stochastic sine circle map as

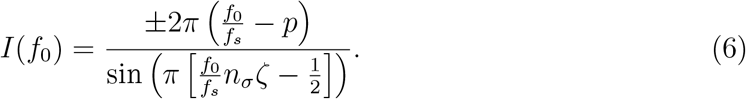

Under the assumption that the standard deviation of the noise is small (*ζ* ≪ 1), the width of the p:1 tongue can be approximated. This requires finding the function *f*_*0*_ ^+^(*I*) demarcating the right boundary of the tongue and the function *f*_0_^−^(*I*) demarcating the left boundary. At the right boundary of the tongue (*f*_*0*_ ^+^/*f*_*s*_ > *p*), a Taylor expansion of the sine term in equation (6) to second order in *ζ* gives the quadratic equation

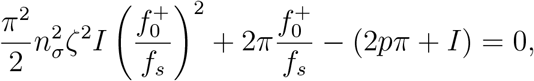

which, after a Taylor expansion to second order in *ζ* of the square root of the quadratic equation discriminant, yields

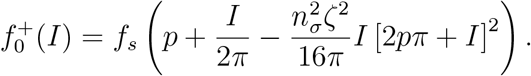

Similarly, we find for the left boundary of the tongue

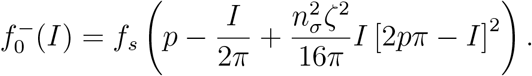

Thus, the width of *p*:1 Arnold tongues can be approximated in the stochastic sine circle map as

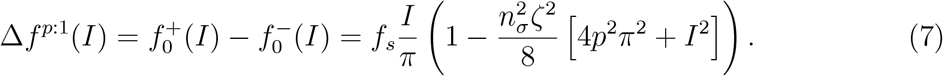

#### 4.1.2 Influence of dithered stimulation on (2*p* − 1):2 Arnold tongues

In the sine circle map, Arnold tongues of order (2*p* − 1):2, *p* ≥ 1, are the second largest after tongues of order *p*:1 (Fig 3A1). Their widths can be approximated by adapting the derivation presented in the previous section. In the deterministic case, (2*p* − 1):2 frequency locking is characterised by 2*p* − 1 complete rotations of the oscillator for every two stimulation cycles. Since two stimulation cycles are considered, (2*p* − 1):2 frequency locking corresponds to stable fixed points of the map given by equation (1) iterated twice, i.e. *θ*_*n*+2_ = *F* (*F* (*θ*_*n*_)), and *θ*_*n*+2_ = *θ*_*n*_ + 2(2*p* − 1)*π*. The random jump size between stimulation pulse *n* and *n* + 2 is

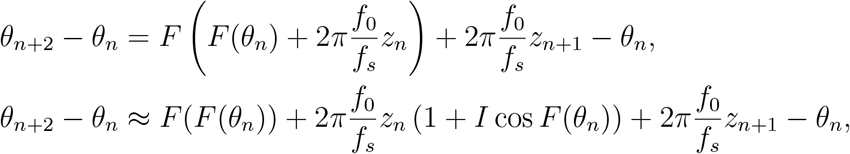

where we have used a Taylor expansion to first order (assuming that *ζ* ≪ 1). Assuming *I* ≪ 1, and since *θ*_*n*_ ≈ *F* (*F* (*θ*_*n*_)) mod 2*π* in the vicinity of the stable fixed points, we have *θ*_*n*+2_−*θ*_*n*_ ≈ 2*π*(*f*_0_/*f*_*s*_) (*z*_*n*_ + *z*_*n*+1_) mod 2*π*. Therefore the jump size is approximately distributed according to 𝒩 (0, *σ*) with 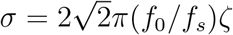.

As before, the shortest trap size *h* is obtained by solving for the position of the stable and unstable fixed points of the deterministic map (see Fig 7C-D for the 1:2 and 3:2 case, respectively). Iterating the deterministic map twice gives

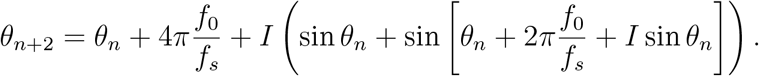

If *f*_0_ is close to the center of the (2*p* − 1):2 tongue, *δ* = (*f*_0_/*f*_*s*_) − (2*p* − 1)/2 is small. Assuming *I* ≪ 1 and *δ* ≪ *I* yields

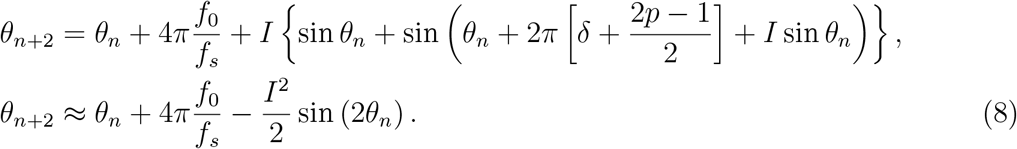

Thus, when (2*p* − 1):2 frequency locking occurs, there are four fixed points which satisfy

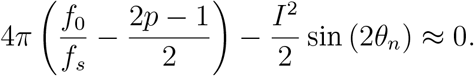

We note that when using equation (8), since the distances between each stable fixed point and the nearest unstable fixed point are the same (see Fig 7C-D), it does not matter at which stable fixed point locking occurs. The trap size *h* is therefore obtained by selecting the shortest distance between one of the stable fixed points and an unstable fixed point. We have for the (2*p* − 1):2 case

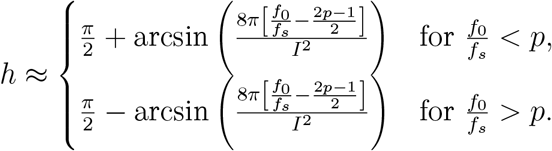

Boundaries for (2*p* − 1):2 tongues in the stochastic sine circle map are obtained from the condition *h* = *n*_*σ*_*σ*, which translates to

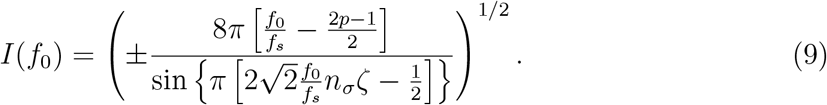

As in the *p*:1 case, the width of (2*p* − 1):2 tongues can be approximated by inverting equation (9) to find the functions *f*_0_^+^(*I*) at the right boundary, and *f*_*0*_^−^(*I*) at the left boundary. At the right boundary of the tongue (*f*_0_^+^/*f*_*s*_ > (2*p* − 1)/2), a Taylor expansion of the sine term in equation (9) to second order in *ζ* give the quadratic equation

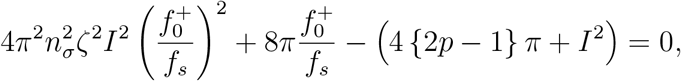

which, after a Taylor expansion to second order in *ζ* of the square root of the quadratic equation discriminant, yields

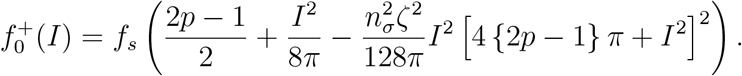

Similarly, we find for the left boundary of the tongue

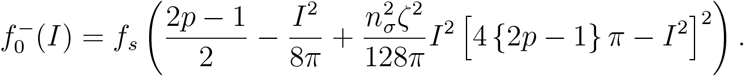

Thus, the width of (2*p* − 1):2 Arnold tongues can be approximated in the stochastic sine circle map as

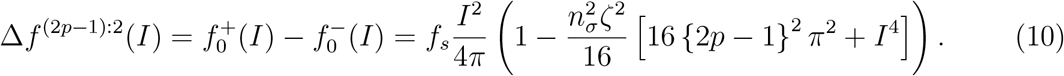

### 4.2 Entrainment metrics used in simulations

We compute several entrainment metrics to characterise the frequency locking behavior of neural oscillators in simulations. Our primary entrainment metric, the rotation number *R*, is measured as

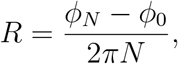

where *N* is large, and *φ*_*n*_ is specified depending on the model considered. In simulations of the sine circle map, we take *φ*_*n*_ = *θ*_*n*_. In simulations of the Kuramoto model, we use *φ*_*n*_ = *ψ*(*t*_*n*_) where *ψ*(*t*) is the phase of the order parameter *Ƶ*(*t*) (see definitions in Section 4.3), and *t*_*n*_ is the time one time step before stimulation pulse *n*. For reference, theoretical definitions of the rotation number in the presence of stochasticity are given in [59, 52].

To detect frequency locking in the presence of noise, the following criterion is used for both models. The system is considered to be entrained to stimulation with a *p*:*q* rotation number if |*R* − *p*/*q*| < tol, and |*S* (***∂****R*/***∂****f*_0_)| < tol′, where *S* (***∂****R*/***∂****f*_0_) is the smoothed partial derivative of the measured rotation number with respect to *f*_0_, and tol and tol′ are tolerances on the rotation number and its smoothed derivative, respectively. These conditions correspond to plateaux in Fig 2 where *p*:*q* frequency locking occurs. For simulations of the sine circle map, we take tol = 6.10^−4^, tol′ = 1.10^−2^, and ***∂****R*/***∂****f*_0_ is smoothed using locally weighted scatterplot smoothing (LOWESS, based on a linear model) with a span of 0.058 Hz (4 samples). For simulations in the Kuramoto model, we take tol = 3.10^−2^, tol′ = 2.10^−2^, and ***∂****R*/***∂****f*_0_ is smoothed using LOWESS with a span of 0.64 Hz (4 samples). These values take into account the different resolutions of the rotation number field for both models in the natural frequency/stimulation amplitude space, simulation duration, and simulation repeats, and were found to robustly detect frequency locking plateaux.

For both models, the width of the *p*:*q* tongue for a particular stimulation amplitude and dithering level is simply measured in simulations as the sum of the frequency width of the discretised bins in the natural frequency/stimulation amplitude space where *p*:*q* entrainment is detected at this stimulation amplitude and dithering level.

Additionally, the mean instantaneous frequency in the Kuramoto model is computed as

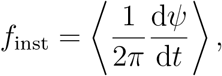

where *ψ* is the phase of the order parameter (definitions in Section 4.3), and ⟨.⟩ denotes the time average over the duration of stimulation.

### 4.3 Simulating populations of coupled neural oscillators

In order to test dithered stimulation in populations of coupled neural oscillators, we simulate *M* = 100 coupled Kuramoto oscillators with noise and homogeneous coupling. The time evolution of the phase *ϕ*_*k*_ of the *k*^th^ oscillator is described by the stochastic differential equation

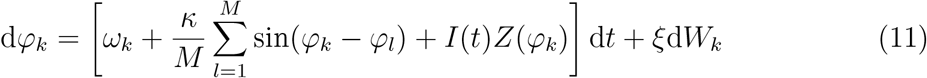

where *ω*_*k*_ is the intrinsic frequency of the *k*^th^ oscillator, *κ* is the coupling strength, *I*(*t*) is the stimulation pulse train, *Z* is the oscillator PRC, *ξ* the model noise standard deviation, and *W*_*k*_ are independent Wiener processes. The order parameter of the network reads 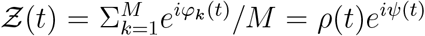 We simulate a more computationally efficient form of equation (11), involving the modulus *ρ* and phase *ψ* of the order parameter, and given by

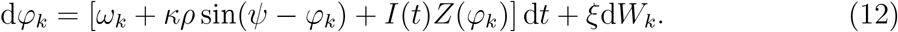

We reproduce signals with dynamics similar to neural oscillations in the absence of stimulation by choosing *κ* = 350, *ξ* = 7.9, and sampling the *ω*_*k*_’s from a Lorentzian distribution centered on the frequency considered (*f*_0_/[2*π*]) and of width 20 Hz. Examples of output signals *χ* (*t*) = ℜ (*Ƶ*(*t*)) [44], where ℜ denotes the real part, are shown in Fig 12A in the supplementary information for two values of *f*_0_. We take *Z* to be the PRC of the standard Hodgkin-Huxley neuron model [48, 49], see Fig 12B1 in the supplementary information.

In Section 2.2, the dithered stimulation pulse train *I*(*t*) is constructed with its stimulation period changing at each stimulation period and given by (1+*z*)/*f*_*s*_, where *f*_*s*_ = 130 Hz is the base stimulation frequency, and *z* is a normal random number sampled from 𝒩 (0, *ζ*^2^) at each stimulation period. In Section 2.3, the stimulation pulse train is constructed as described therein using a finite set of stimulation frequencies. In both cases, contrary to the sine circle map, we can consider square stimulation pulses with a temporal extent, as shown in Fig 12B2 in the supplementary information. The positive stimulation pulse is taken to be 20% of the stimulation period. To avoid harming the brain, charge balance is also enforced, and a negative stimulation pulse occupies the rest of the stimulation period. The magnitude of the positive component is chosen so that the time integral of the stimulation waveform over a period is zero. The model is forward simulated using a Euler-Maruyama scheme with a time step of 10^−4^ s.

## Acknowledgements

We are grateful to Christian Bick for helpful discussions. We would like to acknowledge the use of the University of Oxford Advanced Research Computing (ARC) facility in carrying out this work http://dx.doi.org/10.5281/zenodo.22558

## Data availability statement

No data were collected as part of this work.

## 5 Supplementary information

### 5.1 Phase locking value

The PLV is calculated in the Kuramoto model as

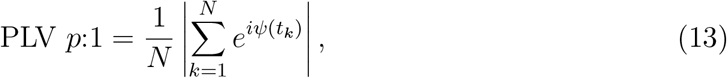

and quantifies the phase concentration of the order parameter at the time of stimulation, considering all stimulation pulses. PLV *p*:1 will therefore detect *p*:1 entrainment, for any *p* ≥ 1. We also consider

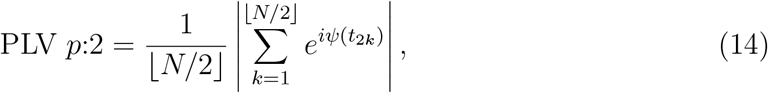

with ⌊. ⌋the floor function. PLV *p*:2 only considers every other stimulation pulse, and will detect *p*:2 entrainment (*p* ≥ 1), which includes for example 1:2 and 3:2 entrainment, but also 1:1 and 2:1 entrainment. Thus, to only detect (2*p* − 1):2 entrainment (in particular the 1:2 and 3:2 tongues), we introduce PLV (2*p* − 1):2 = PLV *p*:2 − PLV *p*:1.

### 5.2 Supplementary figures

**Figure 8:**
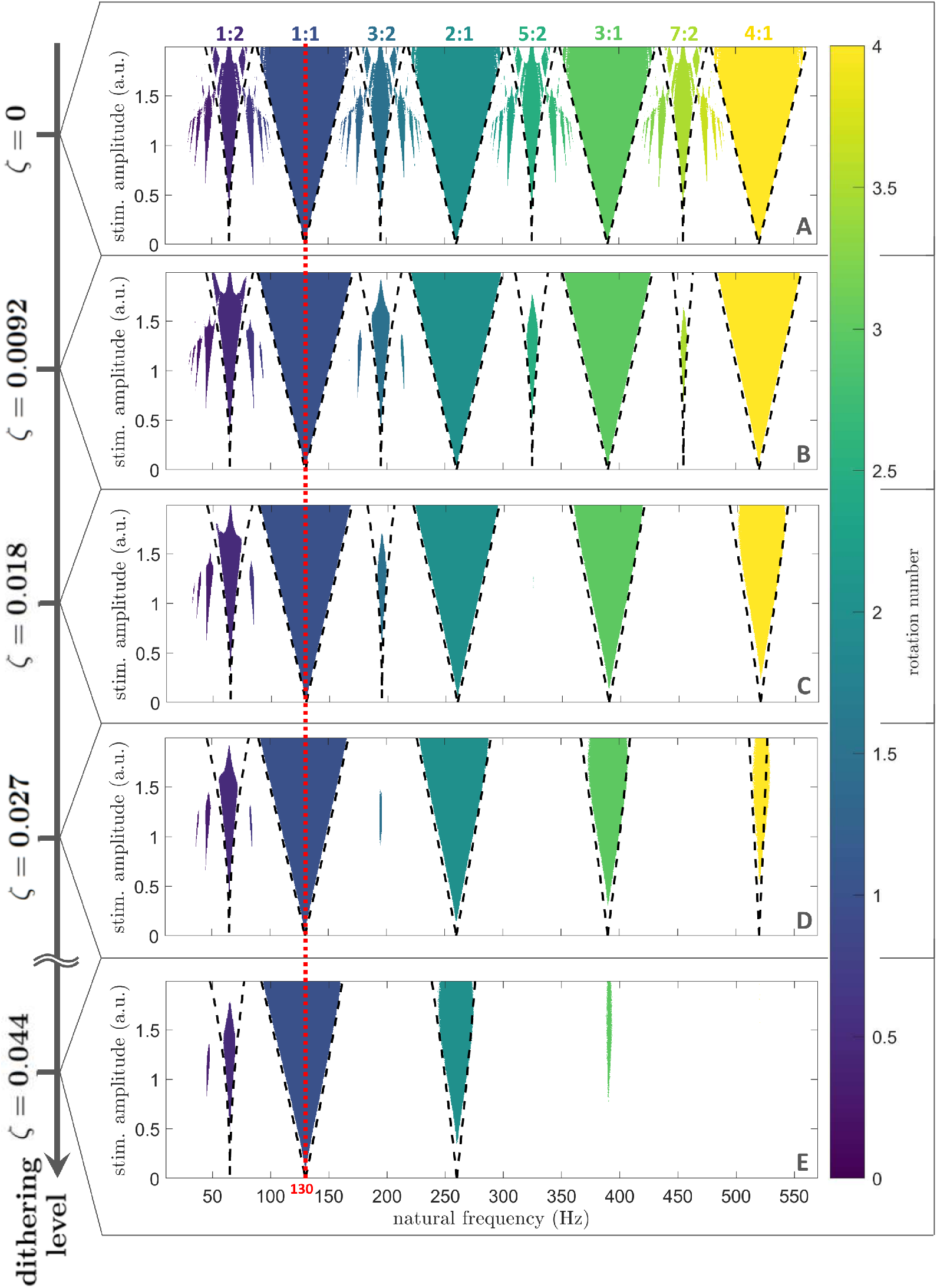
*p*:1 and (2*p* − 1):2 Arnold tongues with *p* up to 4: validation of theoretical results and the efficacy of dithered stimulation. Frequency locking regions in the oscillator frequency/stimulation amplitude space. Only regions of frequency-locking (determined as presented in Section 4.2), are shown in color. The color scale represents the rotation number. Dithering level increases from top to bottom, and theoretical tongue boundaries (equations (6) and (9)) are shown by black dashed lines. The stimulation frequency is indicated by a red dashed line. An animation with finer steps in dithering levels is provided as supplementary material (S2.mp4, see S2 for caption). For each natural frequency, stimulation amplitude, and dithering value, the rotation number is averaged over 10 repeats, with 10^4^ stimulation pulses per repeat. Higher dithering levels, resulting in only the 1:1 tongue being stable, are shown in Fig 3A.

**Figure 9:**
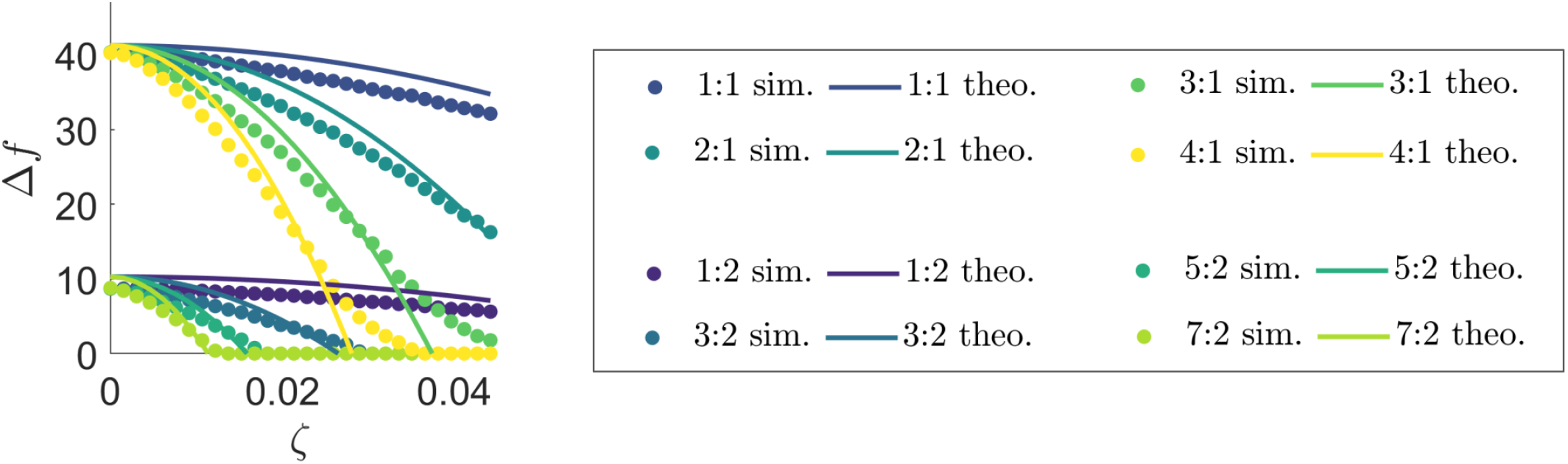
Width of *p*:1 and (2*p* − 1):2 Arnold tongues with *p* up to 4: comparison of simulations and theoretical results. Comparing tongue width obtained from equations (7)/(10) and simulations, as a function of the dithering level, for *I* = 1. For each natural frequency and dithering value, the rotation number from simulations is averaged over 10 repeats, with 10^4^ stimulation pulses per repeat. Higher dithering levels, resulting in only the 1:1 tongue being stable, are shown in Fig 3C2.

**Figure 10:**
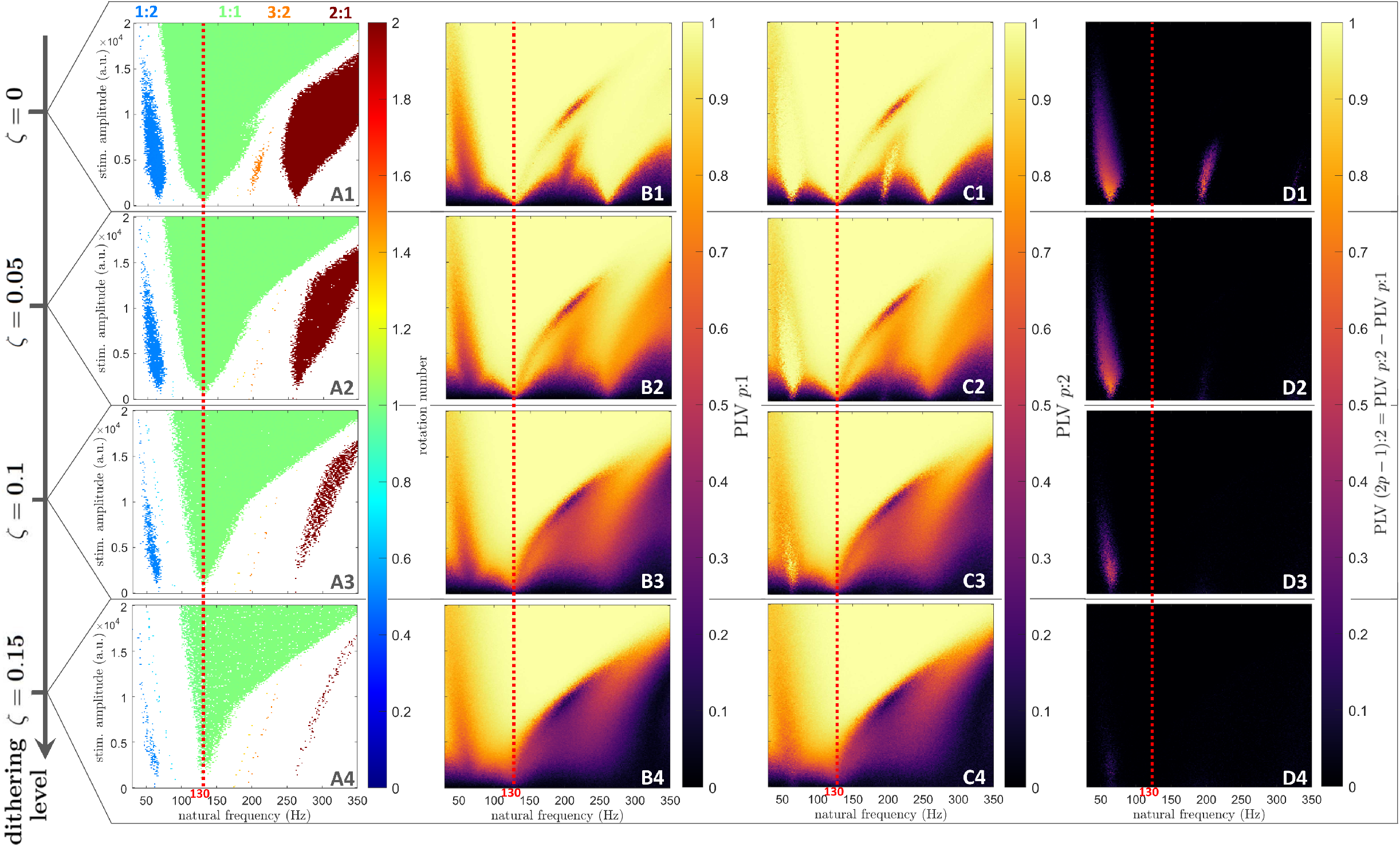
The efficacy of dithered stimulation is confirmed by PLV analysis in populations of coupled neural oscillators. Dithering level increases from the top row to the bottom row. **A:** Frequency locking regions in the natural frequency/stimulation amplitude space. Only regions of frequency-locking (determined as presented in Section 4.2), are shown in color. The color scale represents the rotation number. **B:** PLV *p*:1 (color scale) in the natural frequency/stimulation amplitude space. Here, PLV *p*:1 detects 1:1 and 2:1 entrainment. **C:** PLV *p*:2 (color scale) in the natural frequency/stimulation amplitude space. Here, PLV *p*:2 detects 1:2 and 3:2 entrainment, but also 1:1 and 2:1 entrainment. **D:** PLV (2*p* − 1):2 (color scale) in the natural frequency/stimulation amplitude space, obtained as PLV *p*:2 - PLV *p*:1. Here, PLV (2*p* − 1):2 detects 1:2 and 3:2 entrainment. In all panels, for each natural frequency, stimulation amplitude, and dithering value, the rotation number or mean instantaneous frequency is averaged over 5 repeats, with 400 stimulation pulses per repeat. The stimulation frequency is indicated by red dashed lines. For *ζ* = 0.15, 2:1 entrainment has disappeared while 1:1 entrainment is still supported (compare **B4** to **B1**). 1:2 and 3:2 entrainment have also disappeared (compare **D4** to **D1**). See Section 5.1 for more details on the PLV.

**Figure 11:**
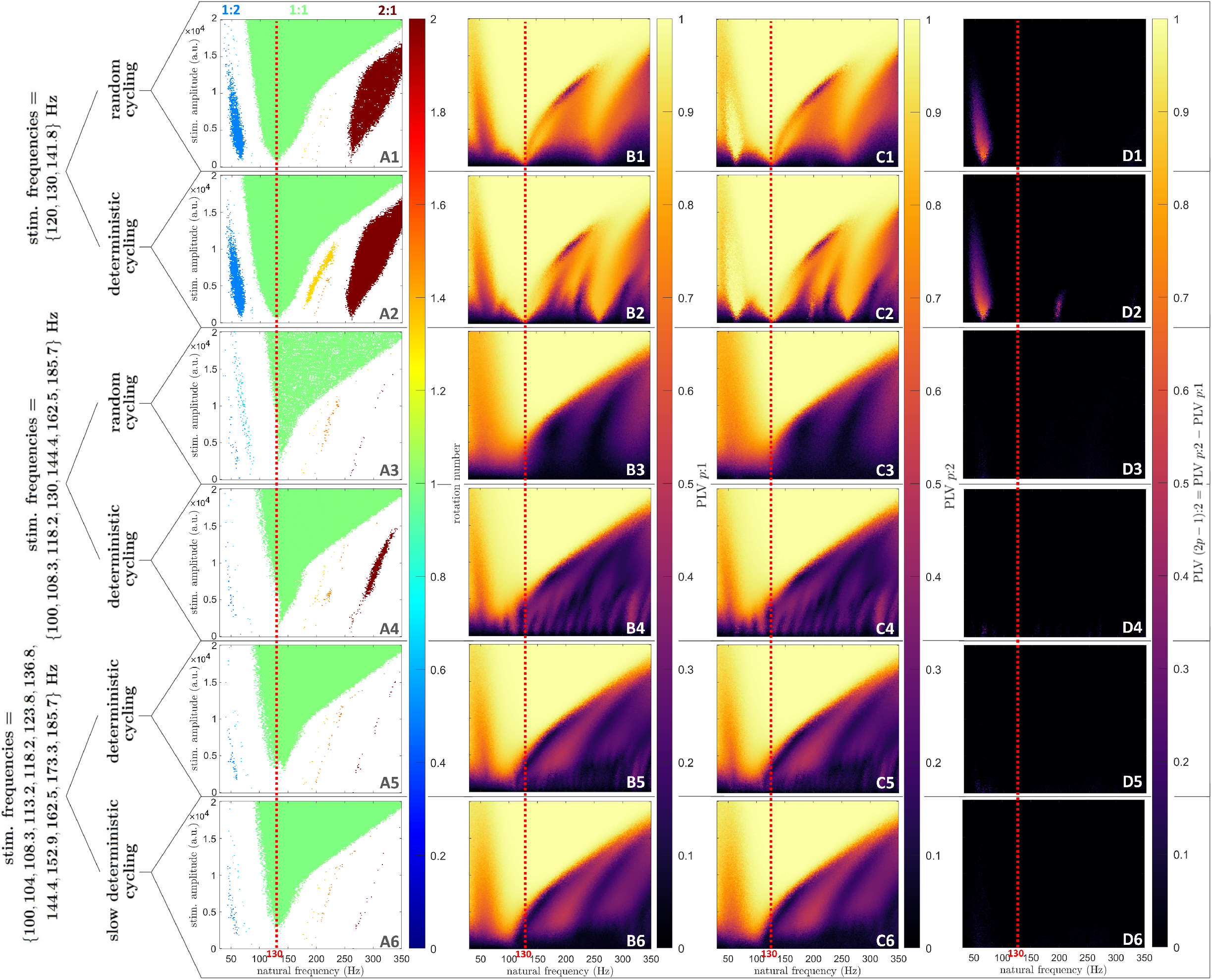
The efficacy of the stimulation frequency toggling approach is confirmed by PLV analysis in populations of coupled neural oscillators. Each row correspond to a particular type of pulse train, as indicated on the left of the figure. For slow deterministic cycling (last row), *N*_*r*_ = 3. **A:** Frequency locking regions in the natural frequency/stimulation amplitude space. Only regions of frequency-locking (determined as presented in Section 4.2), are shown in color. The color scale represents the rotation number. **B:** PLV *p*:1 (color scale) in the natural frequency/stimulation amplitude space. Here, PLV *p*:1 detects 1:1 and 2:1 entrainment. **C:** PLV *p*:2 (color scale) in the natural frequency/stimulation amplitude space. Here, PLV *p*:2 detects 1:2 and 3:2 entrainment, but also 1:1 and 2:1 entrainment. **D:** PLV (2*p* − 1):2 (color scale) in the natural frequency/stimulation amplitude space, obtained as PLV *p*:2 - PLV *p*:1. Here, PLV (2*p* − 1):2 detects 1:2 and 3:2 entrainment. In all panels, for each natural frequency, stimulation amplitude, and dithering value, the rotation number or mean instantaneous frequency is averaged over 5 repeats, with 400 stimulation pulses per repeat. The stimulation frequency is indicated by red dashed lines. See Section 5.1 for more details on the PLV.

**Figure 12:**
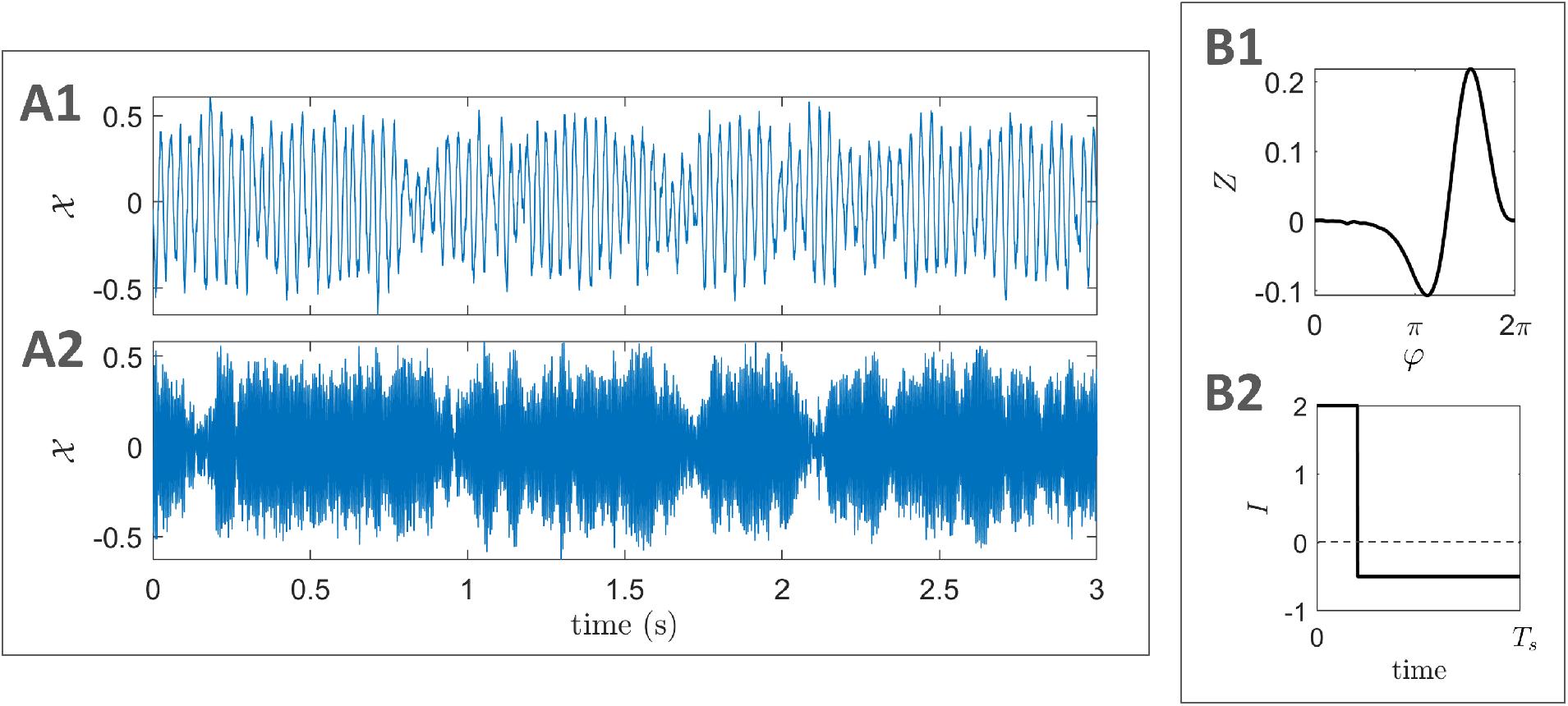
Kuramoto model outputs in the absence of stimulation; PRC and stimulation pulse. Example outputs from the Kuramoto model for the chosen parameters in the absence of stimulation for *f*_0_ = 30 Hz **(A1)** and *f*_0_ = 185 Hz **(A2)**. The PRC of the oscillators is taken from the standard Hodgkin-Huxley neuron model **(B1)**. The charge-balanced rectangular stimulation pulse used in simulations is shown in **B2** (*T*_*s*_ is the stimulation period). The pulse shown has an energy of 1 a.u. over *T*_*s*_, and is taken to be the base pulse for a stimulation amplitude of 1 a.u. in simulations.

**S 1: Animation showing that Arnold tongues disappear at lower dithering levels for higher order entrainment than for 1:1 entrainment in uncoupled neural oscillators (S1.mp4). Top:** Frequency locking regions in the oscillator frequency/stimulation amplitude space. Only regions of frequency-locking (determined as presented in Section 4.2), are shown in color. The color scale represents the rotation number. The dithering level (*ζ*) increases with time, and theoretical tongue boundaries (equations (6) and (9)) are shown by black dashed lines. For each natural frequency, stimulation amplitude, and dithering value, the rotation number is averaged over 10 repeats, with 10^4^ stimulation pulses per repeat. **Bottom:** Example train of stimulation triggers for the given dithering level.

**S 2: Animation validating theoretical results and the efficacy of dithered stimulation for *p*:1 and (2*p* − 1):2 Arnold tongues with *p* up to 4 (S2.mp4). Top:** Frequency locking regions in the oscillator frequency/stimulation amplitude space. Only regions of frequency-locking (determined as presented in Section 4.2), are shown in color. The color scale represents the rotation number. The dithering level (*ζ*) increases with time, and theoretical tongue boundaries (equations (6) and (9)) are shown by black dashed lines. For each natural frequency, stimulation amplitude, and dithering value, the rotation number is averaged over 10 repeats, with 10^4^ stimulation pulses per repeat. Higher dithering levels, resulting in only the 1:1 tongue being stable, are shown in S2. **Bottom:** Example train of stimulation triggers for the given dithering level.

